# HLA-II-dependent neuroimmune changes in Group A Streptococcal Necrotizing Fasciitis

**DOI:** 10.1101/2022.07.04.495809

**Authors:** Ganesh Ambigapathy, Abhijit Satpati, Santhosh Mukundan, Kumi Nagamoto-Combs, Colin K. Combs, Suba Nookala

## Abstract

**Introduction:** Streptococcus pyogenes (Group A Streptococcus, GAS) bacteria cause a spectrum of human diseases ranging from self-limiting pharyngitis and mild uncomplicated skin infections (impetigo, erysipelas, cellulitis) to highly morbid and rapidly invasive life-threatening infections such as streptococcal toxic shock syndrome and necrotizing fasciitis (NF). HLA-Class II allelic polymorphisms are linked with differential outcomes and severity of GAS infections. The dysregulated immune response and peripheral cytokine storm elicited due to invasive GAS infections increase the risk for toxic shock and multiple organ failure in genetically susceptible individuals. We hypothesized that while the host immune mediators regulate the immune responses against peripheral GAS infections, these interactions may simultaneously trigger neuropathology and, in some cases, induce persistent alterations in the glial phenotypes. Here we studied the consequences of peripheral GAS skin infection on the brain in an HLA-II transgenic mouse model of GAS NF with and without treatment with an antibiotic, clindamycin (CLN).

**Methods:** Mice expressing the human HLA-II DR3 (DR3) or the HLA-II DR4 (DR4) allele were divided into three groups: i) uninfected controls, ii) subcutaneously infected with a clinical GAS strain isolated from a patient with GAS NF, and iii) GAS infected with CLN treatment (10mg/kg/5 days, intraperitoneal). The groups were monitored for 15 days post-infection. Skin GAS burden and lesion area, splenic and hippocampal mRNA levels of inflammatory markers, and immunohistochemical changes in hippocampal GFAP and Iba-1 immunoreactivity were assessed.

**Results:** Skin GAS burden and hippocampal mRNA levels of inflammatory markers S100A8/A9, IL-1β, IL-33, inflammasome-related caspase-1 (Casp1), and NLRP6 were elevated in infected DR3 but not DR4 mice. The levels of these markers were significantly reduced following CLN treatment in DR3 mice. Although GAS was not detectable in the brain, astrocyte and microglia activation were evident from increased GFAP and Iba-1 mRNA levels respectively, in DR3 and DR4 mice. However, CLN treatment significantly reduced GFAP immunoreactivity in DR3 mice and not DR4 mice.

**Conclusion:** Our data suggest a skin-brain axis during GAS NF demonstrating that peripherally induced pathological conditions regulate neuroimmune and gliotic events, and CLN may attenuate peripheral infection and subsequent neuroimmune changes in an HLA-II-dependent manner.

## Introduction

*Streptococcus pyogenes*, also known as, Group A Streptococcus (GAS) is a gram-positive β-hemolytic bacterium responsible for a spectrum of infections ranging from self-limiting pharyngitis and mild uncomplicated skin infections (impetigo, erysipelas, cellulitis) to highly morbid and rapidly invasive life-threatening infections such as streptococcal toxic shock syndrome (STSS) and necrotizing fasciitis (NF), a dominant subset of necrotizing soft tissue infections (NSTI) [1]. In most cases, prompt intervention strategies including surgical debridement, antibiotics (combination of clindamycin and penicillin), and management by hyperbaric oxygen therapy lead to effective resolution of the infection and improved outcomes. However, antibiotic-resistant GAS strains are emerging causing prolonged and/or repeated GAS infections and presenting a potential threat to public health [2,3]. Further, the intracellular lifestyle of GAS represents hard-to-reach niches and complicates antibiotic access, thereby resulting in treatment failure leading to recurrent episodes induced from endogenous reservoirs [4]. GAS mimics of host proteins emerging from recurrent infections are not uncommon, and can prime individuals for short- and long-term post-streptococcal autoimmune sequelae that include arthritis, glomerulonephritis, guttate psoriasis, acute rheumatic heart fever, and a temporal association with Sydenham’s chorea, with potential prolonged complications including rheumatic heart disease and clinically heterogeneous pediatric autoimmune neuropsychiatric disorders associated with streptococcal infections (PANDAS) [5–10]. A significant risk for seizures has also been reported to be associated with GAS infections [11]. Several host genetic factors affect GAS NF/STSS pathogenesis. Specifically, polymorphisms in host HLA class II molecules directly determine the risk and significantly influence the outcomes and severity of GAS NF/STSS and post-streptococcal sequelae [12].

It has been established that peripheral inflammation induced by bacteria/virus and/or their products contributes to neuroinflammation, neurodegeneration, and related cognitive dysfunction. For instance, systemic exposure to gram-negative bacterial endotoxin, LPS, has been widely used to demonstrate neuroinflammation and associated behavioral symptoms including sickness behavior [13]. Insults in the brain due to trauma, infections, and peripheral inflammatory mediators prime the microglia. While these resident macrophages of the brain maintain quiescence and perform immune surveillance under physiological conditions, they undergo distinct polarization to a pro- or anti-inflammatory phenotype depending on the type of insult [14]. In the case of pro-inflammatory response, mediators released by activated microglia can, in turn, confer an activated state in responding astrocytes, the predominant glia in the brain, indispensable for the maintenance of neurometabolism [15]. Together, these reactive microglia and astrocytes can promote the progression of neurodegeneration via chronic neuroinflammation [16]. Direct evidence for the role of activated microglia and astrocytes and their inflammatory mediators in neuroinflammation and cognitive impairment has been established as a consequence of HIV-1 and Influenza A infections [17–20]. GAS is not distinctively neurotropic, yet, recurrent GAS infections are frequently associated with several neurological dysfunctions such as Tourette’s syndrome, a PANDAS subgroup of tic disorders, attention disorders [21–24], meningitis [25], and seizure risks in some cases [11]. Intriguingly, while post-streptococcal neurological and neuropsychiatric conditions are associated with an autoimmune response due to cross-reactivity between host and GAS proteins (molecular mimicry) [26,27], it is not known whether invasive GAS NF can induce concomitant neuroinflammatory changes and cause long-lasting brain changes that result in behavioral and cognitive dysfunction. Furthermore, there is a lack of information regarding the neuroprotective potential of CLN, well-established as an effective antibiotic for the treatment of invasive GAS infections [28]. Based on the substantial evidence that HLA-II allelic variations influence peripheral immune responses to invasive GAS infections [12,29] and HLA-II expressing cells reshape T-cell immune responses in the brain during peripheral insults [30], we hypothesized that systemic GAS infections mediate neuroinflammatory changes in an HLA-II dependent manner. Utilizing humanized HLA-II DR3 and DR4 transgenic mouse models of GAS NF/STSS that mimic inflammatory responses and outcomes seen in humans, we tested whether skin GAS infection induces neuroimmune changes that could be attenuated by CLN treatment.

## 2. Materials and Methods

### 2.1 Ethics statement

All the animal experiments described in the current study were conducted in strict accordance with the recommendations in the Guide for the Care and Use of Laboratory Animals of the National Institutes of Health. Breeding, maintenance of mice, and all experimental protocols were approved by the University of North Dakota Institutional Animal Care and Use Committee, protocols 1608-7C and 1704-3.

### 2.2 *In vivo* GAS infections

Male and female mice expressing HLA-II DRB1*0301 (DR3) or DRB1*0401 alleles (DR4) were used. DR3 and DR4 mice were originally generated in the laboratories of Drs. David Bradley, at the University of North Dakota and C.S. David, Mayo Clinic, Rochester, MN [31,32]. Surface expression of HLA-II DR was confirmed by flow cytometry using an LSR-II flow cytometer (BD Biosciences) after staining whole blood with allophycocyanin-labeled anti-HLA-DR antibody (Clone L243) (eBioscience-Thermo Fisher, Waltham, MA) or Tonbo Bio, CA, USA).

A clinical GAS strain (M1 GAS 2006) originally isolated from an NSTI patient (INFECT Consortium) [33,34] was used for *in vivo* studies in the HLA-II transgenic mice. Bacteria growth, preparation of mice, and infections were performed as described previously [29,35,36]. Briefly, GAS 2006 was cultured under static conditions at 37°C for 17 h in THY medium (Todd-Hewitt broth (Bacto BD, Cat# 249240) containing 1.5% (w/v) yeast extract) (Bacto Cat# 212750). The bacteria were centrifuged for 10 min at 1800 rpm (610 x *g*), washed three times, and re-suspended in sterile endotoxin-free Dulbecco’s phosphate-buffered saline (DPBS) (Fisher Scientific, Waltham, MA). GAS bacteria were further diluted to the desired optical density at 600nm (OD_600_ adjusted to yield ∼1-5×10^8^ CFU/0.1mL). Actual inocula were determined by plating serial dilutions on sheep blood agar plates (Thermo Fisher, Waltham, MA). Age-and sex-matched 20–24-week-old, HLA-II DR3 or DR4 mice (n=3-6 mice per group) were used. Since CLN is strongly recommended as the first line of treatment for NSTI [28], we chose CLN (Gold Biotechnology, MO, USA) for the treatment of GAS infections in our mouse models of GAS NF. Seventy-two hours post-infection, mice within each strain group were randomly assigned to receive either treatment with CLN administered intraperitoneally (IP) at 10mg/kg body weight (in 100µL) daily for 5 days or DPBS. Uninfected control mice also received DPBS. Mice were monitored twice a day for 15 days for survival and skin lesion area was measured using digital calipers [29,35].

### 2.3 Tissue collection

At the end of the experiment mice were euthanized by CO_2_ inhalation and blood was drawn through cardiac puncture for bacteremia estimations. The spleen, brain, and necrotic skin were recovered from each mouse under sterile conditions. The necrotic skin was homogenized using a motorized homogenizer (Omni International, Marietta, GA), and the GAS burden was enumerated by preparing tenfold dilutions in DPBS and plated on sheep blood agar as described previously [29,36]. The brains were hemisected, and the hippocampus was isolated from the right hemisphere. The right hippocampus and the spleen were stored in RNA later (Invitrogen, Thermo Fisher, Waltham, MA) for gene expression studies. The left hemispheres were fixed in 4% paraformaldehyde for histological analyses.

### 2.4 Gene expression changes in the spleen and hippocampus by quantitative real-time PCR

Total RNA from the spleen and hippocampal samples were isolated using an RNeasy Mini Kit (Qiagen, Germantown, MD) following the manufacturer’s protocols. RNA concentrations were analyzed using a Nanodrop (ND-1000, Thermo Fisher, Waltham, MA). A total of 0.2-1 µg of RNA was pre-treated with DNAse and used for cDNA synthesis using the iScript cDNA synthesis kit (Bio-Rad, Irvine, CA). Quantitative PCR (qPCR) was performed on Bio-Rad CFX 384 Real-Time PCR system using iTaq-SYBR green supermix (Bio-Rad, Irvine, CA) with specific primer sets (Table 1). Relative gene expression was calculated using the comparative 2^−ΔΔ Cq^ method [37]. Data were normalized against a set of 4 reference genes, ribosomal protein L0 (RPL0), ribosomal protein L27 (RPL27), glyceraldehyde 3-phosphate dehydrogenase (GAPDH), and beta-actin (Actb). Fold change was calculated relative to the average of the uninfected controls and data was represented as relative expression. Primer details are provided in Table 1.

**Table 1.**
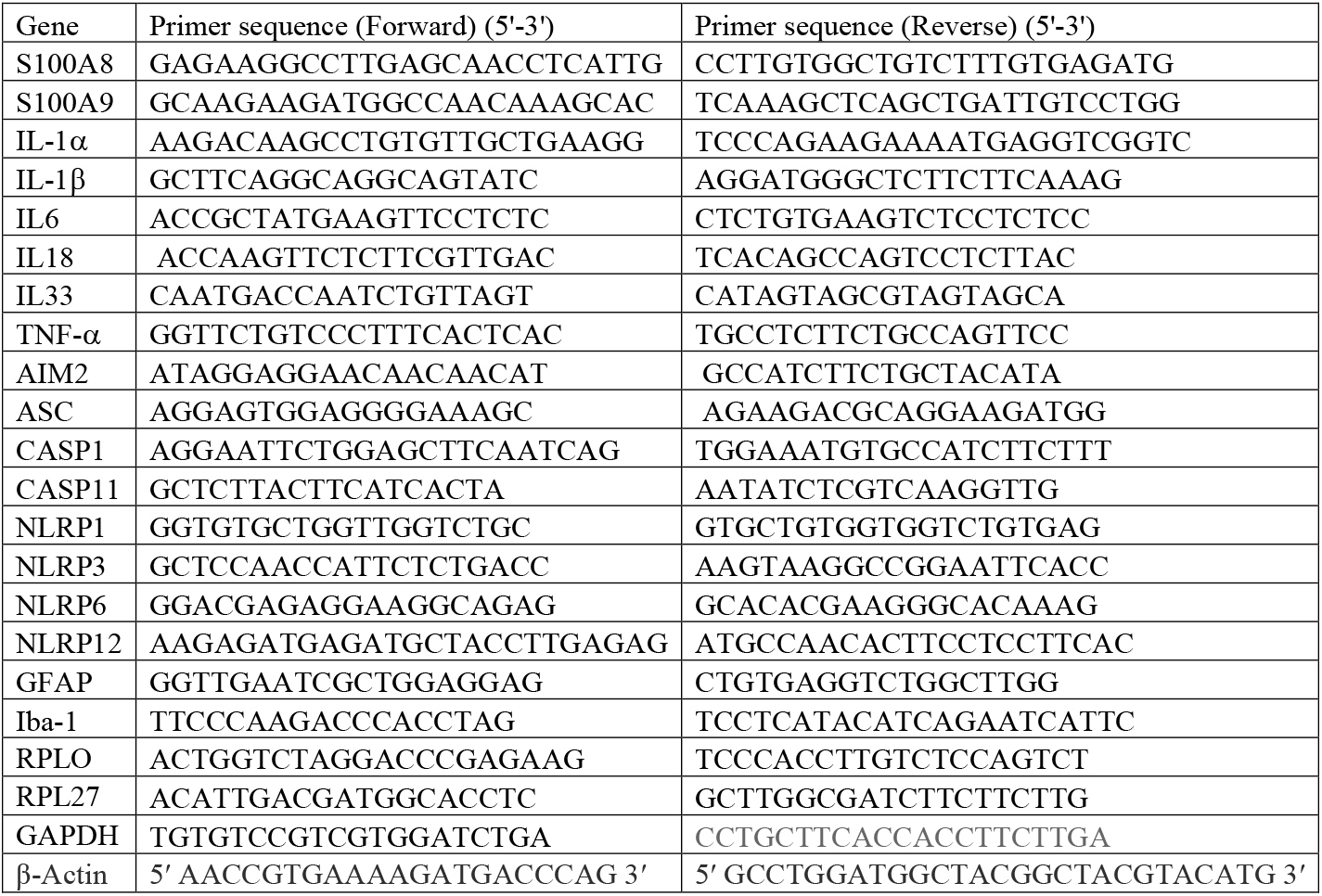
Sequences of primers used for gene expression analysis by quantitative Real Time PCR.

### 2.5 Histological tissue preparation and immunostaining

Paraformaldehyde-fixed brains from uninfected or GAS-infected mice with or without CLN treatment were prepared for immunohistochemical staining as described previously [38]. Briefly, tissues were equilibrated in 30% sucrose prepared in PBS and embedded in a 15% gelatin (in 0.1 M phosphate buffer, pH 7.4) matrix to form a sample block for simultaneous handling of multiple brain samples. The samples were arranged in such a way to facilitate the comparison of different conditions on a single gelatin section. The block was immersed in a 4% paraformaldehyde solution for 3-4 days to fix the gelatin matrix and equilibrated in 30% sucrose change every 3-4 days for 2 weeks during which time the block was completely immersed in the sucrose solution. The cryoprotected blocks were then flash-frozen using dry ice and sectioned at 40 μm using a Leica SM2000R sliding microtome (Leica Biosystems, Deer Park, IL). For immunohistochemistry, sections were first incubated in 0.3% H_2_O_2_ in PBS for 10 min at room temperature to quench endogenous peroxidase activity. Subsequent rinsing, blocking, and antibody incubation was performed with PBS containing 5% normal goat serum, 0.5% BSA, and 0.1% TritonX-100. Anti-ionized calcium-binding adaptor molecule 1 (Iba-1, FUJIFILM Wako Chemicals USA, Richmond, VA) and anti-glial fibrillary acidic protein (GFAP, Cell Signaling Technology, Danvers, MA) antibodies were diluted to 1:1,000 and used to incubate tissue sections overnight at 4 °C. Vectastain ABC Elite Kit (Vector Laboratories, Newark, CA) was used to visualize the immunoreactivity with diaminobenzidine (DAB) as the chromogen according to the manufacturer’s instructions. The sections were mounted onto gelatin-subbed glass slides, cleared in Histo-Clear (National Diagnostics, Atlanta, GA), and cover-slipped using VectaMount (Vector Laboratories).

### 2.6 Quantification of GFAP and Iba-1 immunoreactivity

Immunohistochemically stained slides were digitalized on a Hamamatsu NanoZoomer 2.0-HT slide scanner (Hamamatsu Photonics, Hamamatsu City, Japan) at 40x magnification. The quantification of GFAP and Iba-1 positive cells was performed on whole slide images (n=3-6 mice/group, 2-3 serial sections/mouse), using an open-source digitalized image analysis platform QuPath (v.0.3.2) [39]. For quantification, the image type was set to brightfield H-DAB. To calibrate the intensity of DAB staining and reduce background, areas with distinct positive and negative DAB staining were selected, images preprocessed and RGB values for DAB stain were separated into respective components by applying QuPath’s color deconvolution feature. Using annotation tools in QuPath, the hippocampus region was manually marked as the region of interest for analysis. A thorough manual inspection was performed to exclude any sample that did not exhibit regular boundaries. To consistently capture positive signals that extended beyond the cell body and into the processes (in the case of GFAP), we chose superpixel-based (simple linear iterative clustering, SLIC) segmentation for quantification as described [40–42]. QuPath clusters similar pixels into superpixels based on the RGB values initially set for the DAB stain. We elected to use 25 mm^2^ for the superpixel size to get a precise resolution of the positively stained pixels. Qupath’s built-in intensity feature was then applied to the segmented superpixels to classify them as either “positive” or “negative” involving simple thresholding of a single measurement. Artifacts and blank spaces were selected and ignored from threshold settings and ensuing analysis. The threshold classification was manually checked to avoid false positives or negatives and ensure that the settings captured all positively stained cells in the hippocampus annotated region. Data are presented as a positive % of anti-GFAP and anti-Iba-1-stained pixels.

### 2.7 Generation of heatmaps and correlation plots

Hierarchical cluster heatmaps (Fig. 6A, 6B) and correlation plots (Fig. 6C, 6D) of the expression data from spleen (Sp) and hippocampus (Hc) were generated using Morpheus (Broad Institute; https://software.broadinstitute.org/morpheus). The hierarchal cluster heatmaps were made using the following parameters: one minus spearman rank correlation as the metric, average for linkage method, and clustering by rows and columns. The correlation heatmaps were created using the Morpheus similarity matrix tools, using the following parameters: Spearman rank as the metric, computed for the columns.

### 2.8 Statistical analysis

Values are presented as mean ± SEM and were analyzed using the unpaired Student’s t-test or two-way ANOVA followed by the Sidak post-hoc test or uncorrected Fisher’s LSD under a statistical threshold of P≤.05 using Prism^®^ 9.3.1 (GraphPad Software Inc. San Diego, CA).

## 3. Results

### 3.1 CLN attenuated skin GAS burden in HLA-II DR3 mice

The resolution of inflammation and mitigation of tissue pathology is dependent on both the infecting GAS strain and the host HLA-II context. Here, we investigated the therapeutic benefit of the standard CLN treatment in HLA-II DR3 or DR4 mice. At 15 days post-infection, GAS was not detectable in the blood or brain of untreated or CLN-treated mice (data not shown). GAS persisted in the skin at the site of infection but was significantly reduced with CLN treatment in the HLA-II DR3 mice (*P*=0.0174), but not in the DR4 mice as shown in Fig. 1A. Subcutaneous GAS infections led to the development of lesions at the site of infection that did not change significantly following CLN treatment in DR3 or DR4 mice (Fig. 1B).

**Fig. 1.**
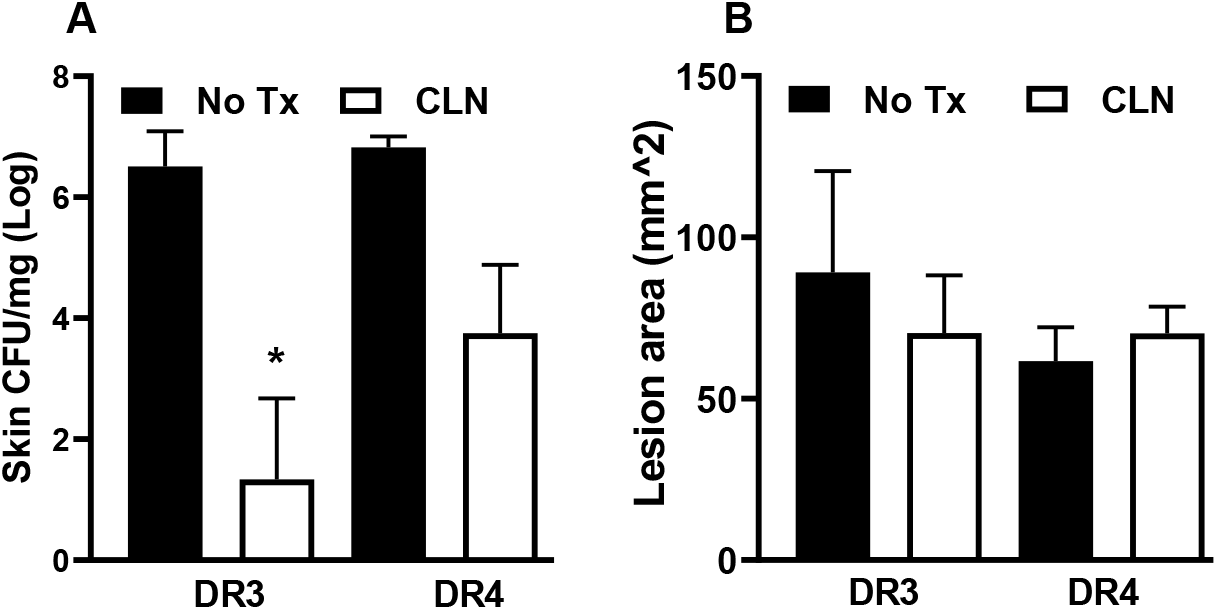
Skin CFU and lesion area in HLA DR3 and DR4 mice following subcutaneous GAS infection. HLA-II mice expressing either DR3 or DR4 were infected subcutaneously with 1 ×10^8^ CFU of GAS 2006. Mice were either untreated (No Tx) or treated with Clindamycin (CLN). At 15 days post-infection, lesions were excised for the enumeration of GAS burden (**A**) and lesion area (**B**). Data are presented as mean values ± SEM, n=3-6, \**P*<.05, Student’s t-test.

### 3.2 CLN treatment differentially altered mRNA levels of numerous genes in spleens from GAS-infected HLA-II DR3 mice

The prolonged skin GAS burden and unmitigated tissue pathology led us to next investigate the changes in splenic mRNA levels of genes associated with inflammation, inflammasome, and pro-inflammatory mediators. Specifically, we examined the changes in the mRNA levels of a) inflammatory markers S100A8, and S100A9; b) inflammasomes NLRP1, NLRP3, NLRP6, and NLRP12, and inflammasome components inflammasome absent in melanoma (AIM2), apoptosis-associated speck-like protein containing a CAR domain (ASC), and caspase 1(Casp1) and caspase 11 (Casp11); and c) proinflammatory mediators IL-1α, IL-1β, IL-6, IL-18, IL-33, and TNF-α. As shown in Fig. 2A, GAS infections in DR3 mice induced a marked increase in the mRNA levels of S100A8 and S100A9 that were significantly reduced (S100A8, *P*=.0104, and S100A9, *P*=.0485) with CLN treatment. Elevated levels of S100A8 and S100A9, mainly derived from neutrophils and macrophages, have been implicated in inducing inflammasome activation and secretion of proinflammatory mediators[43]. Our data show that consistent with the decrease in S100A8 and S100A9 mRNA levels in CLN-treated DR3 mice, mRNA levels of the inflammasomes NLRP1, NLRP3, NLRP6, and NLRP12 also decreased, with a significant reduction in NLRP3 and NLRP6 (*P*=.009 and .011, respectively). Surprisingly, there were significant increases in the mRNA levels of the inflammasome component genes, AIM2 (*P*=.0367), ASC (*P*=.0158), and Casp11 (*P*=.041) in CLN-treated DR3 mice (Fig. 2B). Increases in Casp1 mRNA levels in CLN-treated DR3 mice compared to untreated mice were apparent but these differences did not reach statistical significance (Fig. 2B). Intriguingly, CLN treatment did not significantly alter the mRNA levels of the inflammatory mediators, IL-1α, IL-1β, IL-6, IL-18, IL-33, or TNF-α in the DR3 mice (Fig. 2C). Despite comparable skin GAS burden and lesion area in GAS-infected DR3 and DR4 mice, GAS infections in DR4 mice did not induce increases in the splenic S100A8 or S100A9 mRNA expression (Fig. 2D). Further, except for NLRP1, CLN treatment in DR4 mice did not significantly alter the mRNA expression of other inflammasome markers (NLRP1, *P*=.008, Fig. 2E), or the pro-inflammatory mediators (Fig. 2F).

**Fig. 2.**
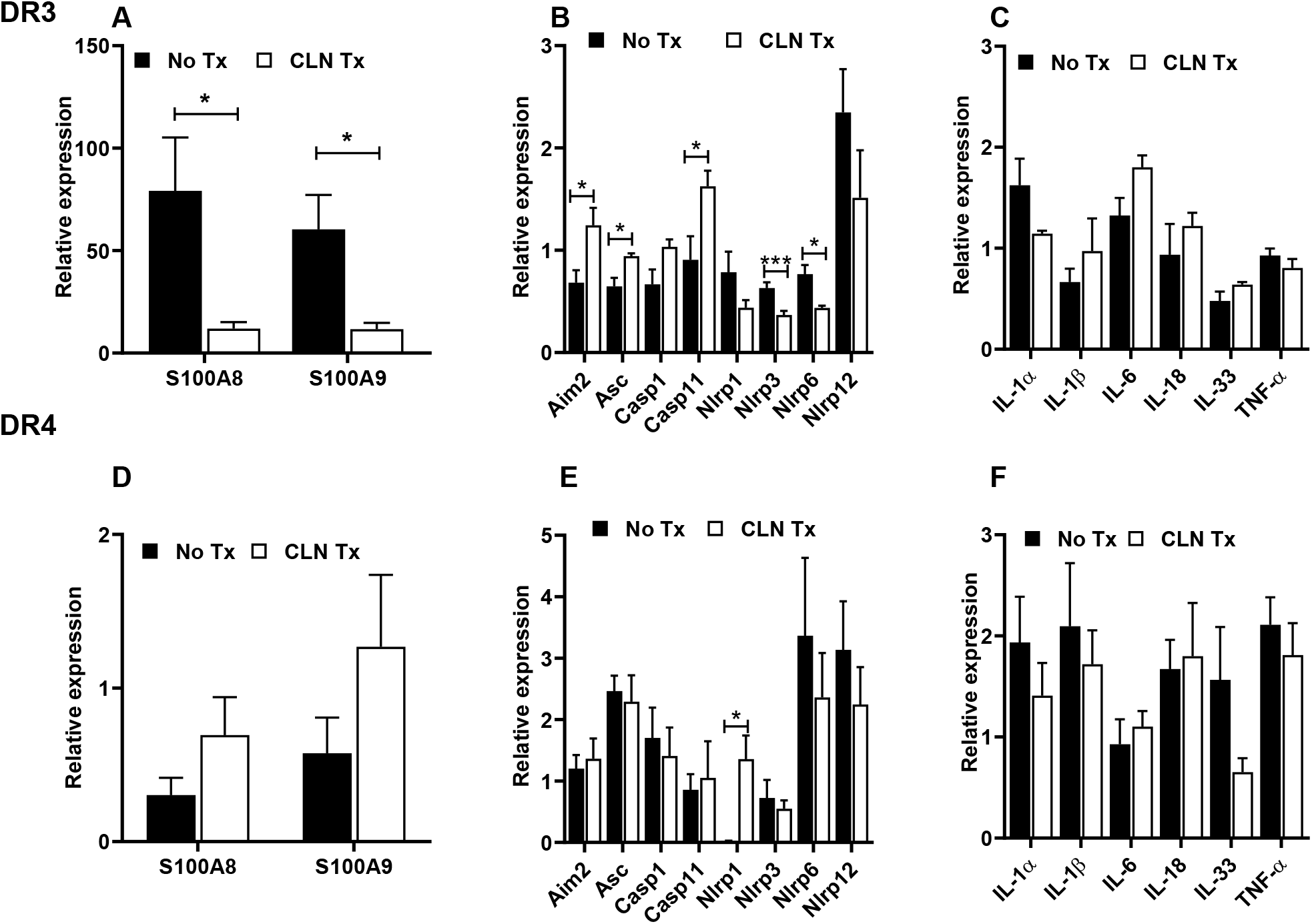
Evaluation of pro-inflammatory markers in the spleen following subcutaneous GAS infection and CLN treatment. The mRNA levels of S100A8 and S100A9 (**A, D**), inflammasome-related genes (**B, E**), and pro-inflammatory mediators (**C, F**) were determined by quantitative real-time PCR in the spleen from GAS-infected HLA-II DR3 (A-C) or DR4 (D-F) mice that were either untreated (No Tx) or treated with CLN. Fold change was calculated using the comparative 2^−ΔΔCq^ method after normalization against a set of 4 housekeeping genes. Data are presented as mean values ± SEM, n=4, * *P*<.05, \*\*\**P* <.001, multiple unpaired t-test.

### 3.3 CLN treatment differentially altered mRNA levels of numerous genes in hippocampi from GAS-infected HLA-II DR3 and DR4 mice

To further examine potential communication of the inflammatory responses to the brain we assessed changes in hippocampal mRNA levels of genes associated with inflammation, inflammasome, and pro-inflammatory mediators as described above. CLN treatment significantly reduced hippocampal S100A9 mRNA levels in the GAS-infected DR3 mice (*P*=.040, Fig. 3A) and to a lesser extent the mRNA levels of S100A8 (*P*=.054, Fig. 3A). However, except for NLRP6 (*P*=.002) and Casp1 (*P*=.034), CLN treatment did not significantly alter the mRNA expression of the hippocampal inflammasomes in GAS-infected DR3 mice (Fig. 3B). Interestingly, mRNA expression of hippocampal IL-1β and IL-33 were significantly reduced in CLN-treated DR3 mice (*P*=.015, and *P*=.006 respectively, Fig. 3C). There was a modest induction of S100A8 and S100A9 mRNA expression in GAS-infected DR4 mice, however, CLN treatment did not ameliorate these responses (Fig. 3D). The relative mRNA expression of NLRP1, NLRP3, NLRP12, and Casp11 showed an increase in CLN-treated DR4 mice, however, increases in NLRP12 levels alone were significant (*P*=.045, Fig. 3E). CLN treatment in DR4 mice did not significantly alter the mRNA expression of the pro-inflammatory mediators (Fig. 3F).

**Fig. 3.**
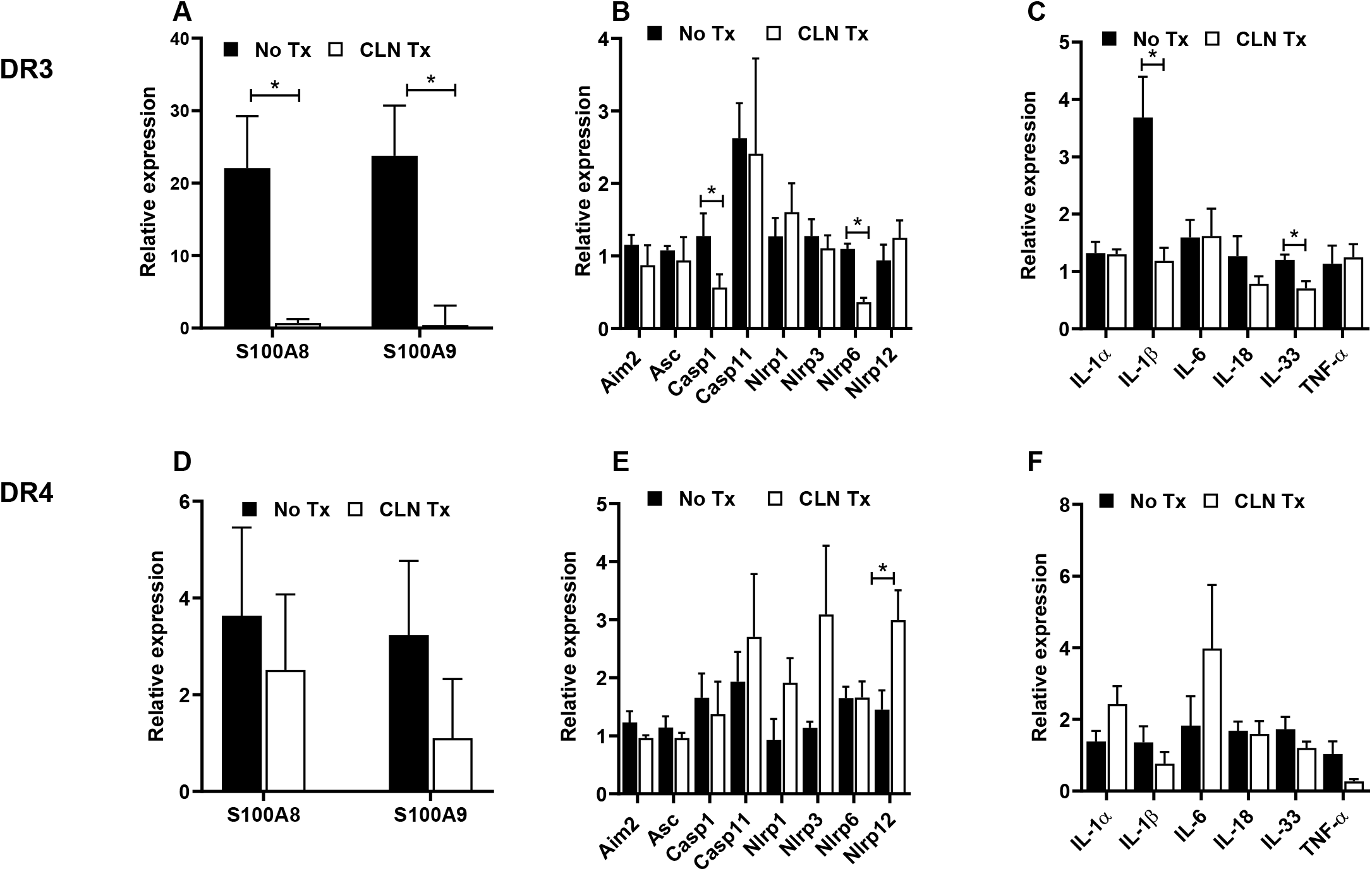
Distinct differences in hippocampal pro-inflammatory and inflammasome markers following subcutaneous GAS infection and CLN treatment. The mRNA levels of S100A8 and S100A9 (**A, D**), inflammasome-related genes (**B, E**), and pro-inflammatory mediators (**C, F**) were determined by quantitative real-time PCR in the hippocampus from GAS-infected HLA-II DR3 (A-C) or DR4 (D-F) mice that were either untreated (No Tx) or treated with CLN. Fold change was calculated using the comparative 2^−ΔΔCq^ method after normalization against a set of 4 reference genes. Data are presented as mean values ± SEM, n=4, * *P*<.05, multiple unpaired t-test.

### 3.4 CLN treatment attenuated GFAP mRNA levels in hippocampi from GAS-infected HLA-II DR3 mice

In the central nervous system, the microglia along with the astroglia play a very important role in brain/hippocampal innate immune responses. Therefore, hippocampal glial activation patterns following peripheral skin GAS infection were studied by quantifying mRNA levels of GFAP and Iba-1 for astrocytes and microglia, respectively, without or with CLN treatment. As shown in Fig. 4A, CLN treatment significantly reduced mRNA levels of GFAP in DR3 mice (*P*=.013) while no such change was observed in DR4 mice (Fig. 4A). Interestingly, Iba-1 mRNA levels were unaltered by CLN treatment in either the DR3 or DR4 mice (Fig. 4B).

**Fig. 4.**
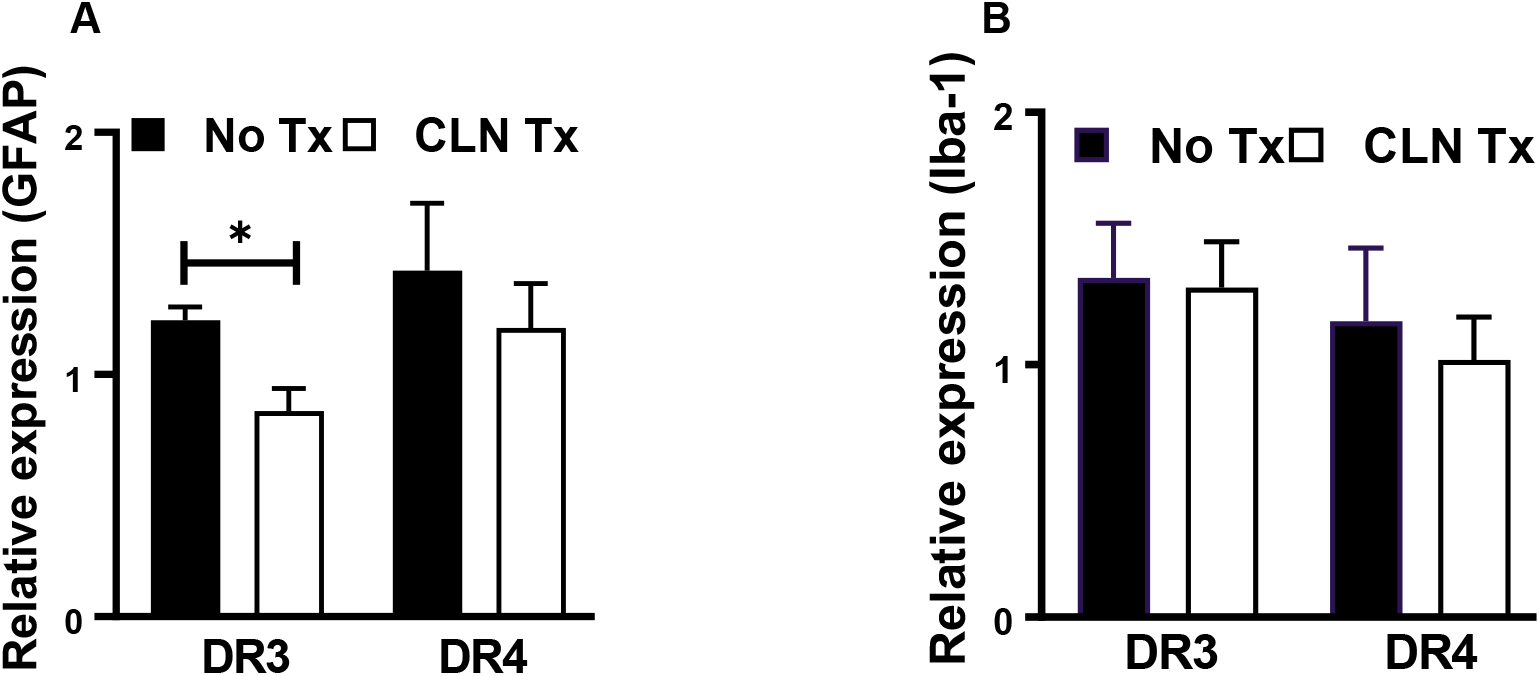
Glial activation as a marker of hippocampal inflammation following subcutaneous GAS infection and CLN treatment. GFAP (**A**) or Iba-1 (**B**) mRNA levels were determined by quantitative real-time PCR in the hippocampi from GAS-infected HLA-II DR3 or DR4 mice that were either untreated (No Tx) or treated with CLN. Bar graphs represent fold change calculated using the comparative 2^−ΔΔCq^ method after normalization against a set of 4 housekeeping genes and uninfected samples. Data are presented as mean values ± SEM, n=4, ** *P*<.05, multiple unpaired t-test.

### 3.5 CLN treatment attenuated GFAP immunoreactivity in hippocampi of GAS-infected HLA-II DR3 mice

We next examined the immunohistochemical changes in GFAP and Iba-1 in DR3 mice to compare to the mRNA changes. Immunohistochemistry demonstrated that while skin GAS infection significantly increased hippocampal GFAP staining (*P*=.002), CLN treatment did not significantly ameliorate this increase (*P*=.203, Fig. 5). Consistent with the mRNA levels, Iba-1 immunoreactivity was unaltered in DR3 mice (Fig. 5).

**Fig. 5.**
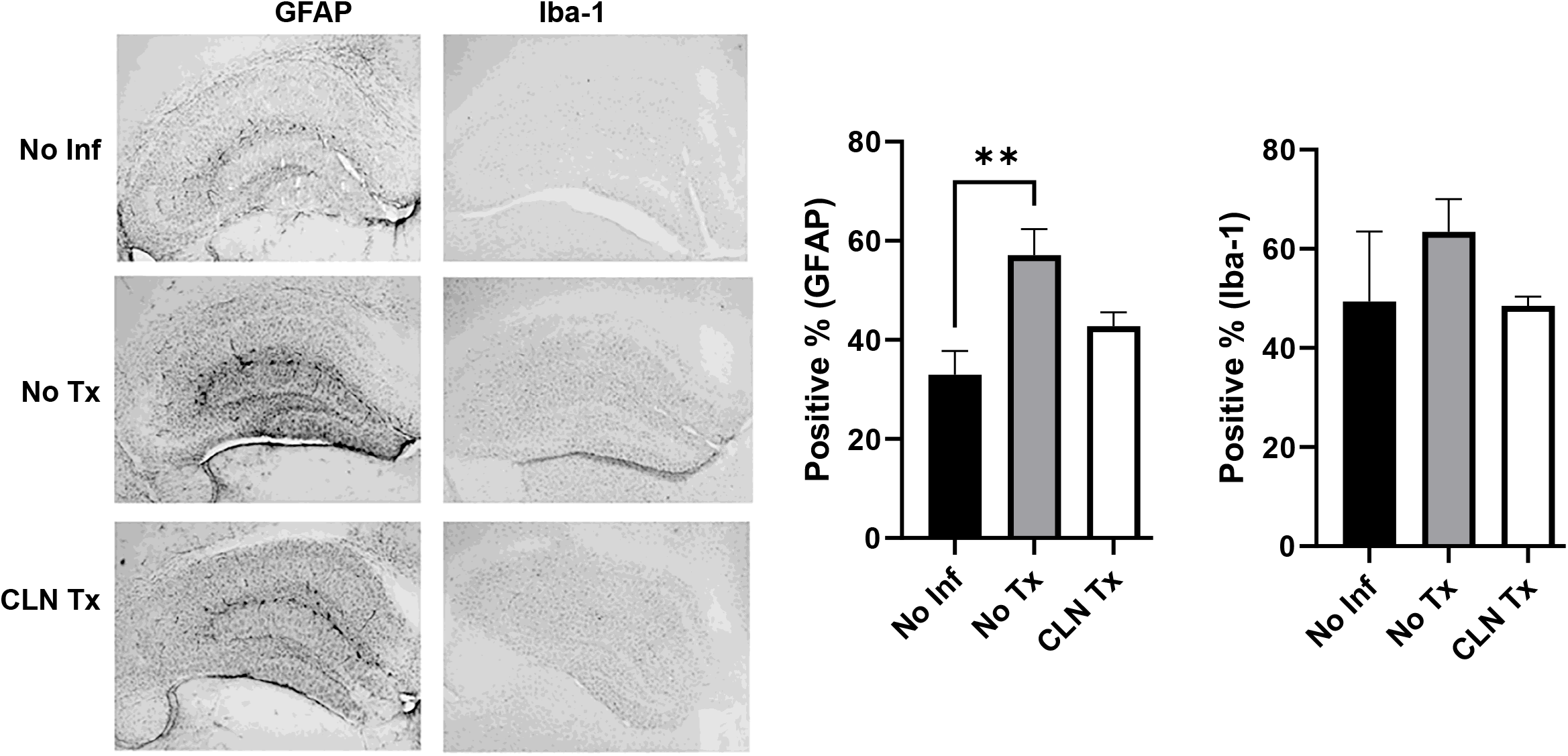
Clindamycin treatment attenuated GFAP immunoreactivity in hippocampi of GAS-infected HLA-II DR3 mice. Representative image of immunohistochemical detection of the astrocyte marker, GFAP, and the microglial marker, Iba-1 in the brain tissue from Uninfected (No Inf) or GAS infected -/+ CLN Tx HLA-II DR3 mice collected 15 days post-infection. GFAP and Iba-1 intensity were quantitated using QuPath. Bar graphs represent the positive % of stained pixels calculated using Qupath software. Data are presented as mean values ± SEM, n=3-4, Student’s t-test.

### 3.6 Analysis of mRNA expression patterns by heat map and similarity matrix revealed unique clusters

To further discern the transcriptional regulation of splenic and hippocampal S100A8, S100A9, inflammasome, and pro-inflammatory mediators in skin GAS infections, we performed a more in-depth analysis based on clusters or similarity index of the mRNA expression profile of the markers. The relative expression values in mRNA levels for the S100A8, S100A9, inflammasomes, and pro-inflammatory mediators were uploaded to MORPHEUS software to generate heat maps and a similarity matrix. As shown in Fig. 6A, three main clusters were apparent in the heatmap of the spleen: cluster-1 made up of AIM2 and IL-18; cluster-2 made up of ASC, Casp1, IL-1β, IL-33, NLRP6, TNF-α, IL-1α, NLRP12, and NLRP3; and cluster-3 made up of NLRP1, Casp11, IL-6, S100A8, and S100A9. Clusters in hippocampus included: cluster-1 Casp1, IL-18, AIM2, ASC, GFAP and IL-33; cluster-2 made up of Casp11, NLRP1, IL-1α, NLRP6, NLRP3, IL-6, and NLRP12; and cluster-3 made up of TNF-α, IL-1β, S100A9, S100A8, and Iba-1 (Fig 6B). It is notable from Spearman similarity analysis that the expression of spleen S100A8 and S100A9 coordinated with IL-6, Casp11, and NLRP1 (Fig 6C), while hippocampal S100A8 and S100A9 coordinated with Iba-1 and the two most prominent pro-inflammatory mediators TNF-α and IL1-β (Fig 6D).

**Fig. 6.**
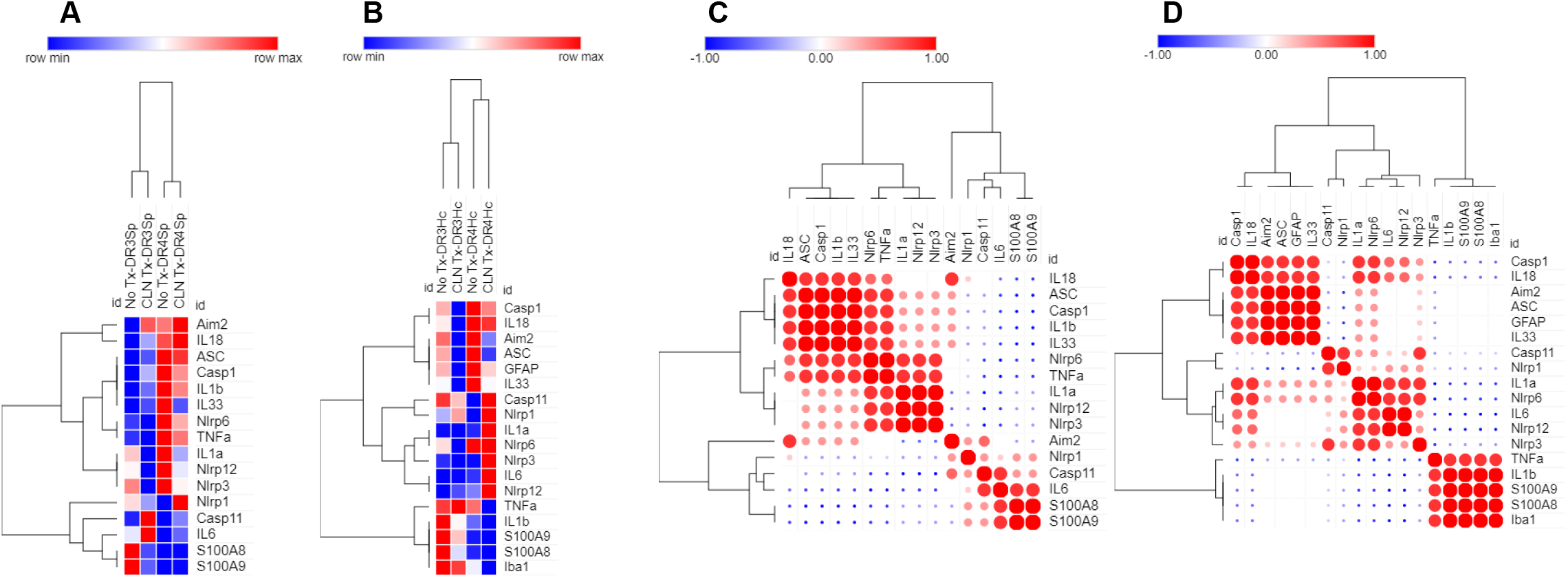
Generation of heatmaps and correlation plots. Hierarchical cluster heatmaps and correlation plots of the expression data from the spleen (Fig. 6A, 6C) and hippocampus (Hc, Fig. 6B, Fig. 6D) were generated using Morpheus (Broad Institute; https://software.broadinstitute.org/morpheus). The hierarchal cluster heatmaps were made using the following parameters: one minus spearman rank correlation as the metric, average for linkage method, and clustering by rows and columns. The correlation heatmaps were created using the Morpheus similarity matrix tools, using the following parameters: Spearman rank as the metric, computed for the columns.

## Discussion

The severity of GAS infection outcomes depends on the heterogeneity of the GAS virulence factors as well as the host HLA-II allelic polymorphisms. GAS infections have been linked to a spectrum of neurological complications arising as direct effects of autoimmune reactions [27] as well as indirect effects from peripheral inflammation[44]. Dileepan et al. have demonstrated that intranasal GAS infections induce CNS complications, blood-brain barrier compromise, IgG deposition, microglial activation, and infiltration of a GAS-specific Th17 subset of CD4+ cells, despite the lack of viable GAS persistence in the brain tissue in C57BL/6, C57BL/6J, or SJL/J female mice [45]. To establish the relevance of our findings to human settings, we used humanized mice expressing HLA-II DR3 or DR4, as preclinical subcutaneous infection models of GAS NSTI with no apparent lethal systemic toxicity [12,29,46]. We assessed the effectiveness of CLN, a protein synthesis inhibitor antibiotic, with a wide spectrum of activities including activity against stationary growth-phase GAS and the suppression of virulence GAS toxins [47], that has been demonstrated in Swiss Webster and C57BL/6 mouse models of GAS myositis and subcutaneous infections [2,48,49]. In parallel with the findings reported by Andreoni et al [2], our results show that CLN treatment does not eliminate skin GAS burden but likely attenuates the activity of GAS virulence factors.

It is well established that the calgranulin S100A8/A9 complex is one of the biomarkers in sepsis and exerts its inflammatory role through TLR4 activation [50]. Consistent with the preferential reduction in the skin GAS burden in CLN-treated DR3 mice, elevated mRNA levels of splenic and hippocampal S100A8 and S100A9 were significantly reduced in DR3 but not DR4 mice. The reasons underlying the differential induction of S100A8 and S100A9 responses in GAS-infected DR3 and DR4 mice are not clear and need further investigations. It will be important to assess the changes in the virulence capability of GAS bacteria in different hosts that might influence inflammation and outcomes. The disparity in CLN efficacy is also concerning and adds to the overriding effects of HLA-II allelic polymorphisms in shaping not just peripheral but also brain inflammatory responses during skin GAS infections.

In the present study, hippocampal mRNA levels of Casp11 and IL-1β were highly induced in GAS-infected DR3 mice suggesting the possibility that despite the lack of viable GAS burden in the brain, GAS products or GAS genetic material likely triggered these responses. The functional role of AIM2 inflammasome and caspases in regulating astrogliosis has been reported [51,52]. In support of this notion, it is interesting to note from our similarity matrix that changes in the relative expression of GFAP showed positive coordination with AIM2 inflammasome, and associated partners ASC, Casp1, IL-33, and IL-18. Ma et al. demonstrated that the AIM2 inflammasome negatively regulates microglial activation in mouse models of EAE [53]. The involvement of AIM2 in regulating microglia responses in GAS NF is unclear. In our study, skin GAS infection-induced mRNA expression of AIM2, ASC, and Iba-1 in DR3 and DR4 mice, and their levels were not ameliorated with CLN in DR3 or DR4 mice. Monocytes and macrophages are the main sources of pro-inflammatory mediators IL1-β, IL-18, TNF-α, and IL-6 which are also central to GAS pathogenesis. TNF-α is mainly released by activated microglia and it is well known that the innate immune mechanisms and inflammasome signaling are mediated by microglia in the brain. Our data show the mRNA levels of hippocampal TNF-α, IL-6, and Iba-1 persisted despite CLN treatment in DR3 mice. In contrast, hippocampal mRNA levels of both TNF-α and Iba-1 levels showed a modest decrease in CLN-treated DR4 mice. Increased hippocampal NLRP1 and NLRP12 mRNA expression was an unpredicted outcome of CLN treatment in GAS-infected DR3 and DR4 mice. It has been shown that the NLRP1 inflammasome is expressed by pyramidal neurons and oligodendrocytes in the brain [54], and increased NLRP1 expression has been reported in aging-related neuronal damage[55]. Among the inflammasomes, the cytosolic pathogen sensor, NLRP12, has been implicated in the maintenance of gut homeostasis and is different from other members of the NLR family due to its dual role in activation and dampening of NF-κB signaling[56]. Whether increased hippocampal NLRP1 and NLRP12 expression is an unintended consequence of CLN therapy causing gut dysbiosis and alterations in brain inflammasomes through the gut-brain axis is an open critical question. Further studies are needed to elucidate the involvement of hippocampal NLRP1 and NLRP12 in regulating glial and neuronal responses in GAS infections.

We acknowledge the limitations of our study including the small sample size. However, this proof of concept study demonstrates HLA-II-dependent neuroinflammation despite CLN therapy and might impactfully translate to other disease entities. With respect to the potential underlying mechanisms, in addition to HLA-II allelic variations, the likelihood of neuroinflammatory sequelae may be influenced by GAS strain variability. Therefore, one future direction is to evaluate the neuroinflammation potential of several clinical strains of GAS across a battery of HLA-II transgenic mice. We considered HLA-II transgenic mice as a clinically relevant and translational model to study the neuroinflammation effects of peripheral inflammation involved in GAS infections since these mice mimic responses seen in humans [29,57]. Notwithstanding that the magnitude of responses is directly linked to HLA-II allelic variations, it is understandable that a single gene cannot hold the key to all the intricacies of disease pathogenesis underlying GAS infections. Several genes related to an immune function whose loci are located within the central MHC region or in linkage disequilibrium with HLA-II genes might either counteract or exacerbate the overriding effects of HLA-II genes thereby polarizing the immune responses and influencing outcomes [26].

In conclusion, our study adds to the growing evidence linking pro-inflammatory insults and long-lasting pathological changes with behavioral complications and cognitive dysfunction. Our findings indicate that subcutaneous GAS infections that display systemic inflammation trigger the production of pro-inflammatory mediators and glial changes, despite the absence of viable GAS burden in the brain, raising the possibility of increased risk of neurological changes after invasive GAS infections, especially among those individuals who are genetically “at-risk”.

## Statement of Ethics

This study protocol was reviewed and approved by the University of North Dakota Institutional Animal Care and Use Committee, protocols 1608-7C and 1704-3.

## Conflict of Interest Statement

The authors have no conflicts of interest to declare.

## Funding Sources

The work presented was supported by the grants from the European Union (FP7/2012–2017) under the grant agreement 305340, grants from the Swedish Research Council under grant number 20150338, UND CoBRE Host-Pathogen Interactions Pilot Award supported by the NIH/NIGMS award P20GM113123, and UND VPRED Post-Doctoral support to A. Satpati. Histological services were provided by the UND Histology Core Facility supported by the NIH/NIGMS award P20GM113123, DaCCoTA CTR NIH grant U54GM128729, and UND SMHS funds.

## Author Contributions

Ganesh Ambigapathy, Santhosh Mukundan, and Abhijit Satpati acquired data and analyzed experiments, revised the manuscript, and approved the final manuscript. Santhosh Mukundan., Colin K. Combs, and Kumi Nagamoto-Combs were involved in experiment interpretation, revision of the manuscript, and final approval of the manuscript. Suba Nookala designed all the experiments, acquired and interpreted results, drafted the first version of the manuscript, and approved the final version.

## Data Availability Statement

All data associated with the manuscript are included in the manuscript.

## Notes

### Competing Interest Statement

The authors have declared no competing interest.

